# “Mesenchymal Osr1+ cells regulate embryonic lymphatic vessel formation”

**DOI:** 10.1101/2023.12.11.571189

**Authors:** Pedro Vallecillo-García, Mira Nicola Kühnlein, Mickael Orgeur, Nils Rouven Hansmeier, Georgios Kotsaris, Bernd Timmermann, Claudia Giesecke-Thiel, René Hägerling, Sigmar Stricker

**Affiliations:** Freie Universität Berlin, Institute for Chemistry and Biochemistry, 14195, Berlin, Germany; Department of Hematology, Oncology and Tummorimmunology, Charité-Universitätsmedizin Berlin, Corporate Member of Freie Universität Berlin and Humboldt-Universität zu Berlin, Germany; Institut Pasteur, Université Paris Cité, Unit for Integrated Mycobacterial Pathogenomics, 75015 Paris, France; Research Group ‘Lymphovascular Medicine and Translational 3D-Histopathology’, Institute of Medical and Human Genetics, Charité-Universitätsmedizin Berlin, Augustenburger Platz 1, 13353 Berlin, Germany; Berlin Institute of Health at Charité-Universitätsmedizin Berlin, BIH Center for Regenerative Therapies, Augustenburger Platz 1, 13353 Berlin, Germany; Research Group ‘Development and Disease’, Max Planck Institute for Molecular Genetics, Ihnestraße 63-73, 14195 Berlin, Germany; Berlin Institute of Health at Charité-Universitätsmedizin Berlin, BIH Academy, Clinician Scientist Program, Charitéplatz 1, 10117 Berlin, Germany; Max Planck Institute for Molecular Genetics, 14195, Berlin, Germany

## Abstract

The lymphatic system is formed during embryonic development by the commitment of specialized lymphatic endothelial cells (LECs) and their subsequent assembly in primary lymphatic vessels. While lymphatic cells are in continuous contact with mesenchymal cells during development and in adult tissues, the role of mesenchymal cells in lymphatic vasculature development remains poorly characterized. Here, we show that a subpopulation of mesenchymal cells expressing the transcription factor *Osr1* are in close association with migrating LECs and established lymphatic vessels in mice. Lineage tracing experiments revealed that Osr1+ cells precede LEC arrival during lymphatic vasculature assembly in the back of the embryo. Using Osr1-deficient embryos and functional *in vitro* assays, we show that *Osr1* acts in a non-cell autonomous manner controlling proliferation and early migration of LECs to peripheral tissues. Thereby, mesenchymal Osr1+ cells control in a bimodal manner the production of extracellular matrix scaffold components and signal ligands critical for lymphatic vessels formation.

## Introduction

The lymphatic vasculature establishes a blind-ended hierarchical network of vessels with crucial roles in interstitial fluid homeostasis, immune cell response and lipid metabolism. In peripheral tissues, lymphatic vessels form thin-walled capillaries that drain interstitial fluid and facilitate the transport of macromolecules and cells into larger pre-collecting lymphatic vessels. Here, the lymph is moved unidirectionally by the cooperative action of lymphatic valves, the synchronic contraction of lymphatic vessel-associated mural cells and passively by the force generated in surrounding tissues such as skeletal muscles and arteries. Finally, the lymph is transported into the blood stream at the subclavian vein (1–3).

Lymphatic vessels are formed by specialized endothelial cells, lymphatic endothelial cells (LECs), that originate from venous and non-venous tissues depending on the vascular bed (4–6). By far best understood is the formation of LECs via dedifferentiation of embryonic venous endothelial cells. In mice, reprograming of venous cells towards a LEC fate occurs at the embryonic days (E)9-E10 relying on the activation of key LEC transcription factors such as Prospero Homeobox 1 (PROX1) and SRY-related HMG-box 18 (SOX18), and is associated with the expression of markers as lymphatic vessel endothelial hyaluronan receptor 1 (LYVE1)(1, 2, 7). After this initial step in LEC commitment, which occurs at the dorsal part of the cardinal vein, LECs delaminate and migrate into the surrounding mesenchyme forming transient primordial lymphatic vascular structures called primordial thoracic duct, formerly known as lymph sacs, which can be observed in the mouse embryo during the stages E11.5-E14.5 (1, 2, 8, 9). LEC initial migration from the cardinal vein is dependent on the production of vascular endothelial growth factor C (*Vegfc*) and the activation of its primary receptor fms-related tyrosine kinase 4 (FLT4), also known as VEGFR3 (10, 11). Activation of canonical or non-canonical VEGFR3-signaling, and enzymes controlling the proteolytic activation of VEGFC play an essential role in LEC initial migration as well as development and maintenance of lymphatic vessels (10, 12–15). In addition, directed migration of LECs is controlled by the CXCL12/CXCR4 chemokine axis (16–18). Both ligands, *Vegfc* and *Cxcl12*, are highly expressed by embryonic mesenchymal cells adjacent to the developing lymphatic vasculature (10, 12, 16, 19). Despite the knowledge gained in the mechanism supporting LEC migration in the past years (20), little is known how the final pattern of lymphatic vessels in different tissues of the embryo is achieved. Several cell types have been described to influence lymphatic vessel development in zebra fish and mice, such as platelets, arterial endothelial cells, neurons, myeloid cells and mural cells (12, 16, 21, 22). However, embryonic mesenchymal cells adjacent to lymphatic vessels have remained understudied in part due to the lack of specific markers that label mesenchymal cell subpopulations during development. Of note, mesenchymal cells are important producers of the extracellular matrix (ECM) scaffold during embryogenesis and adult life (23). LEC-ECM interactions and biophysical properties of the ECM associated to lymphatic vessels influence lymphatic vessel development and function (24, 25).

The transcription factor *Osr1* is expressed in a variety of mesenchymal cells derived from the lateral plate and intermediate mesoderm (26, 27). In the limb, Osr1+ mesenchymal cells are present before skeletal muscle progenitors colonize the limb bud mesenchyme, and they produce guidance and a proper ECM scaffold for skeletal muscle patterning (28). *Osr1* expression decreases in late fetal stages of development but is reactivated after tissue damage (19, 28–30). We recently showed that *Osr1* is required in mesenchymal cells to organize lymph node lymphatic vasculature assembly, and that *Osr1*-expressing cells cooperate with LECs in the lymph node anlage driving lymph node initiation (19). This suggests a crosstalk between mesenchymal Osr1+ cells and LECs as an important event for lymphatic vessel formation. Moreover, the lymphatic vasculature can be remodeled in pathogenic conditions such as inflammation, wound healing, tumor formation, hypertension or tissue transplantation (4, 31), where mesenchymal cell-LEC interactions might be necessary to achieve lymphangiogeneis. Despite the important functions assigned to mesenchymal cells, the role of these cells or the ECM produced by them in the formation of murine lymphatic vasculature remain obscure.

Here we show that mesenchymal cells expressing the transcription factor *Osr1* are in close association with the developing venous-derived lymphatic endothelial cells. Functionally, lack of *Osr1* revealed a non-cell autonomous function of embryonic Osr1+ mesenchymal cells that control lymphatic vessel formation by producing a beneficial ECM scaffold and signaling molecules necessary for LEC directed migration and proliferation.

## Methods

### Animals

Mice were maintained in an enclosed, pathogen-free facility, and experiments were performed in accordance with European Union regulations and under permission from the Landesamt für Gesundheit und Soziales (LaGeSo) Berlin, Germany (Permission numbers ZH120, G0346/13, G0240/11, G0268-16). Mouse lines were described previously; *Osr1^GCE^*(26), *R26R^mTmG^* (56), *Osr1^LacZ^* (29), *Cxcr4*^+/-^ (47).

### Tamoxifen and Progesterone administration for Osr1^+^ cell lineage tracing

As we described previously (19, 28), Tamoxifen (Sigma Aldrich) was dissolved in a 1:10 ethanol/sunflower oil mixture. For lineage tracing experiments, we bred *R26R^mTmG^*^/mTmG^ females to *Osr1^GCE^*^/+^ males. Pregnant females were injected with 150 µl of a 20 mg mL^-1^ Tamoxifen stock. Tissues were collected at E14.5.

### Tissue preparation

Embryonic tissues were fixed in 4% PFA for 2h on ice. Tissues were dehydrated in two steps using 15% and 30% (w/v) sucrose (Roth) solutions before O.C.T. (Sakura) cryo-embedding in a chilled ethanol bath. Embryonic tissue was sectioned at 12 or 100 μm thickness.

### Immunolabelling

Cryo-sections were warmed up for at least 30 minutes at room temperature (RT). Sections and E14.5 isolate skin tissue were blocked with 5% (v/v) Horse Serum (Vector Laboratories) in 0.1% (v/v) Triton X-100 (Sigma Aldrich) PBS for 1h at RT. Primary antibodies dissolved in blocking solution were incubated at 4 °C overnight, followed by secondary antibody staining of 1h at RT. Antibodies used for these experiments are listed in supplementary tables 1 and 2. Specimens were counterstained with 5µg µL^-1^ 4’, 6-diamidino-2-phenylindole (DAPI; Invitrogen) and mounted with FluoromountG (SouthernBiotech).

### Cell isolation and flow cytometry

Isolation of E13.5 Osr1+ cells has been described before (19, 28). Briefly, inner organs, tail, limbs and cranial tissue above the tongue were removed from E13.5 *Osr1^GCE^*^/+^ and *Osr1^GCE^*^/GCE^ embryos. Next, embryonic tissue was minced using a small scissor in 1 ml high-glucose Dulbecco’s modified eagle medium (DMEM, Pan Biotech) containing 10% foetal bovine serum (FBS, Pan Biotech) and 1% penicillin/streptomycin (P/S) solution. Further enzymatic digestion was performed using 0.7 mg ml^-1^ of Collagenase (Collagenase A, Roche) in DMEM medium at 37 °C for 45 min. For the isolation of E13.5 LECs and BECs, embryonic tissue was dissected from E13.5 *Osr1^controls^* and *Osr1^GCE/GCE^*embryos and treated as above described for E13.5 Osr1+ mesenchymal cells. Antibody labelling (antibodies see supplementary table 3) was performed for 20 min on ice.

Isolation of primary LECs (tdLECs) was described before (57). Briefly, tails of at least 7 wild type adult animals (8-30 weeks) were used. Tails were cut at the attachment site and washed twice with Hank’s Balanced Salt Solution (HBSS) containing 1% P/S. Next, epidermis and dermis were isolated mechanically from the underlying musculoskeletal system. Isolated tissue was cut in pieces of 2 cm and digested for 1h at 37°C in a HBSS solution containing 1% P/S and 2 U ml^-1^ Dispase II (Roche). Epidermal layer was separated from the dermis using two tweezers. Collected dermal tissues were further digested enzymatically in 30 ml DMEM medium containing 10% FBS, 1% P/S and 1 mg ml^-1^ collagenase A (Roche) for 90 minutes. Digested tissue was filtered through a 100 µm cell strainer and cells were collected by centrifugation at 300g for 10 min. Cell suspension were cultured on 0.4% gelatine coated dishes using LEC medium that contains high-glucose DMEM, 20% FBS, 1% P/S solution, 10 µg/ml endothelial cell growth supplement (ECGS) (Thermo scientific), 50 µM 2-Mercaptoethanol, 1% Non-essential amino acids solution (Thermo scientific) and 50 µM for 4-7 days. Prior FACS isolation, cells were detached using for epitope conservation and washed once with HBSS supplemented with 1% P/S and 0.4% FBS. Antibody labelling was performed for 20 min on ice.

Before flow cytometry, cell suspensions were washed using a solution containing PBS, 0.4% FBS and 2 mM EDTA, collected by centrifugation at 300 g for 5 min and passed through a 35-µm cell strainer filter (BD Biosciences). To assess viability, cells were stained with propidium iodide (2 μg ml^−1^, eBioscience) immediately before sorting or analysis.

Sorts and analyses were performed on a FACS Aria II and FACS Aria fusion (BD Biosciences). Data were collected using FACSDIVA software. Further analyses were performed using FlowJo 10 (FlowJo LLC) software. Sorting gates were defined based on unstained and fluorescence negative controls. Cells were collected into 400 µl high-glucose DMEM containing 10% FBS, and 1% P/S solution for Osr1+ mesenchymal cells or LEC medium.

### Conditioned media and wound healing assay

For the production of conditioned media, 80.000 E13.5 *Osr1^GCE^*^/+^ and *Osr1^GCE^*^/GCE^ cells isolated by FACS were plated in 24 well plates. After 100% confluence was reached, Osr1+ cells were culture in 300 µl high-glucose DMEM containing only 1% P/S solution. After 24 h, media containing secreted molecules from E13.5 *Osr1^GCE^*^/+^ and *Osr1^GCE^*^/GCE^ cells were collected and used as a conditioned medium. Conditioned media from several time points was collected.

For wound healing assays, 24 well plate were coated with 0.4 % gelatine and 20.000 pdLECs (P3-P5) were seeded into two wells culture-inserts (Ibidi®). After 45 min two wells culture-inserts were removed and gap closure was monitorized every 2 hours with a Leica DMi8 microscope.

### ECM deposition and decellularization

For ECM deposition, 80.000 embryonic E13.5 *Osr1^GCE^*^/+^ and *Osr1^GCE^*^/GCE^ cells isolated by FACS were plated in 10 mm coverslip coated with 1 mg/ml fibronectin and cultured in DMEM containing 10% FBS and 1% P/S solution until 100 % confluence. Next, cells were cultured for 3 weeks in DMEM containing 10% FBS, 1% P/S solution and 2 µ/ml ascorbic acid. ECM produced by Osr1+ cells was decellularized using a freeze/thaw method as described previously (28). Briefly, culture medium was aspirated and cultures were washed with PBS. Next, PBS was removed and exchanged by 300 µl distilled water. Cultures were frozen in a -80°C freezer and thawed in a 37°C water bath. Remaining water was careful aspirated by pipetting and tdLECs were seeded on the decellularized ECM (dECM). To assess tdLECs proliferation cultures on dECM, they were cultured for 48 h in LECs medium.

### Imaging

X-Gal staining of whole lymph nodes or the dermis of the ear were documented with a Zeiss SteREO Discovery V12 stereomicroscope. Confocal images of immunolabelled sections were taken using the confocal laser scanning microscope systems LSM710, LSM810 (Zeiss) or Leica DMi8 microscopes. Images were captured using Zen 2010 (Zeiss) and LAS Life System (Leica). For the quantifications shown in figures 2D, 2G, S2F, 6E, 6G, 6H and 7D consecutive images were selected and measurements from a single embryo are depicted in the same colour.

### Quantitative real-time PCR

Total RNA extraction from FACS isolated cells was performed using Direct-zol^TM^ RNA MicroPrep (Zymo Research) following manufactureŕs protocol. Reverse transcription was conducted using the M MuLV Reverse Transcriptase Kit (Biozym). Relative gene expression analyses were performed using GoTaq® qPCR kit (Promega) or Blue ŚGreen qPCR kit (Byozim) on a 7900HT Real Time PCR system or QuantStudio 7 Flex Real-Time-PCR-System (Applied Biosystems). Primer sequence information is provided in supplementary table 4. Data were acquired and analysed using SDS 2.0 and QuantStudio^TM^ Real-Time PCR softwares (Applied Biosystems).

### Transcriptome analysis

To obtain the total RNA amount necessary for RNA-seq, we pulled LECs from 8-10 E13.5 *Osr1^cantrols^* embryos and 8-10 *Osr1^GCE/GCE^*embryos per sample. E13.5 LECs were isolated by FACS as described above. Total RNA was isolated using Direct-zol RNA Microprep (ZYMO RESEARCH) following manufactureŕs protocol. Total RNA was measured using a Qubit 4 Fluorometer and RNA quality was assessed using an Agilent RNA 6000 Nano kit prior library preparation. Library preparation was conducted according to Illumina instructions TruSeq Library Preparation Kit V2. Next, libraries were subjected to high-throughput sequencing using an HiSeq 2500 device. Obtained fastq data were further analysed using the platform Galaxy Europe (https://usegalaxy.eu). 53-81 million of reads were obtained and mapped using STAR (PMID: 23104886) against the genome of *Mus musculus* version mm10. Quantification of aligned reads at the gene level was conducted using featureCounts (PMID: 24227677). Differential gene expression analysis was performed using DESeq2 (PMID: 25516281). Transcript per million (TPM) abundances were calculated using normalised gene counts from DESeq2 analysis of E13.5 *Osr1^GCE/GCE^* LECs samples. Genes with an absolute log2 fold change of at least 0.3, a Benjamini-Hochberg adjusted p-value (padj) below 0.05, and TPM value higher than 1, were considered as being differentially expressed between E13.5 *Osr1^cantrols^*and *Osr1^GCE/GCE^* LECs. Gene ontology (GO) analysis was performed using Enrichr (PMID: 23586463). Ligand-receptor interaction network was drawn using graph-tool v2.45_5 (https://graph-tool.skewed.de/) with a hierarchical edge bundling. Raw fastq and count data were uploaded in the Gene Expression Omnibus (GEO) database under the accession number GEO.

### Statistical analysis

Student’s t-test and one-way ANOVA with Dunnett’s post-hoc comparison were performed using Prism 8 (GraphPad) software. Error bars in all figures, including supplementary information, represent the mean ± standard error of the mean (s.e.m.). In figure 6G, a paired Student’s t-test was used due to the differential proliferation showed by pdLECs isolated from different experiments.

## Results

### Osr1+ mesenchymal cells accompany LECs during development and in adult tissues

Impaired lymphatic vasculature assembly in embryonic lymph node anlage and back lymphedema found in Osr1-deficient embryos indicated a more general role of Osr1+ mesenchymal cells in lymphatic vasculature formation (19, 27). Therefore, we analyzed *Osr1* expression and distribution of Osr1+ cells in association to the developing lymphatic vasculature. Using an Osr1-GFP reporter line (*Osr1^GCE/+^*) (26), we identified Osr1+ cells in close association with LECs during the first stages of lymphatic vessel development. At E11.5, Osr1+ cells were found in the mesenchyme populated by delaminating and migrating PROX1+ LECs dorsally and ventrally from the cardinal vein (figure 1A, 1A’ and S1A). Of note, at E11.5 Osr1+ cells were not found in the vicinity of back skin tissues (figure 1A’’). At E12.5, we observed that the primordial thoracic duct was surrounded by a mesenchyme rich in Osr1+ cells (figure 1B). At E14.5, when the first lymphatic vascular structures are established in the dermis of the back, whole-mount immunofluorescence of E14.5 *Osr1^GCE/+^* isolated skin showed dermal lymphatic vasculature embedded by Osr1+ dermal fibroblasts (figure 1C and S1B).

**Figure 1.**
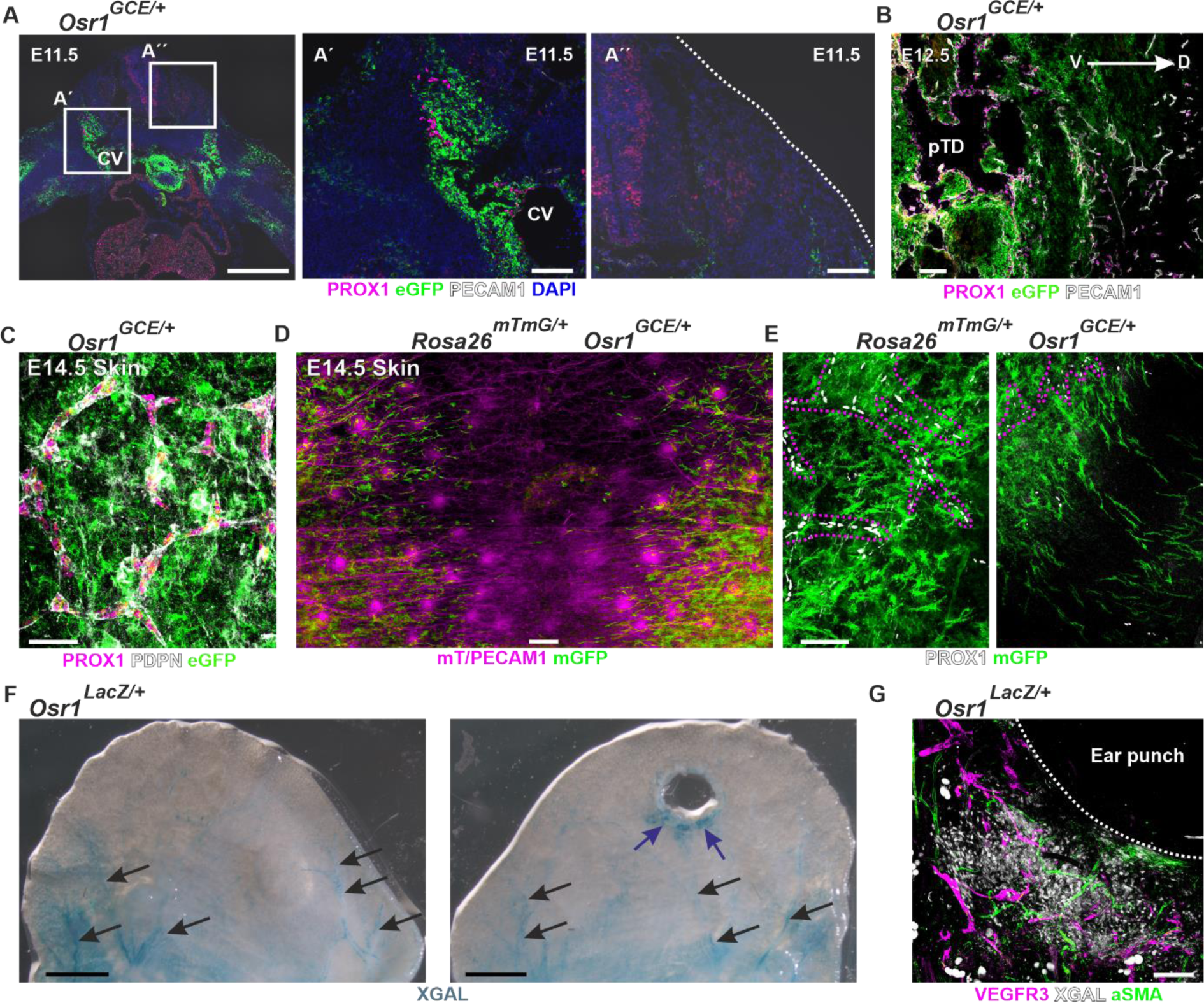
Mesenchymal Osr1+ cells accompany lymphatic endothelial cells during development. **(A)** Immunofluorescence of E11.5 Osr1GCE/+ cross-section shows Osr1+ cells (eGFP) in the migration path of delaminating LECs labeled by PROX1 and PECAM1 dorsally of the cardinal vein. Boxed regions are shown at higher magnification at the right. **(B)** E12.5 sagittal section of Osr1GCE/+ embryos show primordial thoracic duct surrounded by Osr1+ mesenchymal cells. **(C)** E14.5 whole-mount immunofluorescence shows lymphatic vasculature labeled by PDPN and PROX1 embedded in Osr1+ dermal fibroblasts in the skin of E14.5 Osr1GCE/+ embryos. **(D, E)** Whole-mount immunofluorescence of E14.5 Rosa26mTmG/+ Osr1GCE/+ skins shows the distribution of lineage traced Osr1 mesenchymal cells (mGFP). Stripes of Osr1-traced cells are localized ahead of blood vasculature labeled with mT/PECAM1 and lymphatic vasculature labeled with PROX1. Dashed lines represent the border of lymphatic vessels in (E). **(F)** Whole-tissue X-gal staining revealed Osr1 expression in the dermal side of the ear from adult Osr1LacZ reporter animals in contralateral (left) and ear punch-injured (right). Blue arrows point to activated Osr1 expression in the injury area and black arrows to Osr1 expression in association to established vessels. **(G)** Osr1 expression close to the ear punch injury in adult Osr1LacZ reporter animals assessed by whole-tissue X-gal staining. Dashed lines represent the border of the regenerating tissue. Representative immunofluorescence images have been captured from at least 3 different animals. Scale bar represents in **A** 500 µm, **A’** and **A’’** 100 µm, **B** 100 µm and 50 µm (E14.5), **C** 200 µm, **D** 2 mm and **E** 100 µm. In **A**, CV represents cardinal vein. In **B**, V represents ventral, D dorsal and pDT primordial thoracic duct.

In order to follow Osr1+ mesenchymal cell descendants and their contribution to the mesenchyme adjacent to blood and lymphatic vasculature, we performed whole-mount immunofluorescence of E14.5 *Rosa26^mTmG/+^ Osr1^GCE/+^* back skin after tamoxifen induction at E11.5. Genetic tracing of E11.5 Osr1+ mesenchymal cells (schematic depiction in figure S1C) showed contribution to dermal fibroblasts between blood and lymph vessels (figure 1D), to mural smooth muscle actin-expressing cells (αSMA+) in arteries, and to cells associated with veins in the skin (figure S1C). In the back skin, Osr1+ descendants were found in the lymphatic avascular midline suggesting that Osr1+ cells progress ahead of LECs in the back dermis (figure 1E) anticipating LEC migration through the forming dermis. Of note, Osr1+ descendants were located in close association with blood and lymphatic vasculature but in agreement with previous results (19) mGFP signal was not found in endothelial cells (figure S1D).

Analysis of *Osr1* expression in embryonic stages and adult tissues have revealed that *Osr1* expression decrease at late stage of development (28–30) and Osr1+ cells are found in the stroma of some adult tissues (19, 28, 30). We analyzed Osr1+ cell distribution in association with blood and lymphatic vasculature in adult ear dermis and lymph nodes of *Osr1^LacZ/+^*animals. Whole-mount LacZ staining revealed that Osr1+ cells are associated with arteries in the dermal tissue of the ear and to lymphatic vessels in the medulla of mesenteric lymph nodes of *Osr1^LacZ/+^* animals (figure 1F and S1E). Ear punch trauma led to an activation of *Osr1* expression in surrounding mesenchyme (figure 1F, right panel, S1F) in cells closely associated with lymphatic and blood vasculature (figure 1G and S1G).

In summary, Osr1+ cells accompany LEC migration during lymphatic vasculature formation in the embryo and persist as mesenchymal vessel-associated cells including mural cells in adult tissues.

### Osr1+ mesenchymal cells control lymphatic vasculature formation

Since *Osr1*-expressing mesenchymal cells associated with migrating LECs and embedded the first lymphatic vasculature in the skin, we asked whether impairment of the lymphatic vasculature could be responsible for the back edema observed in *Osr1^GCE/GCE^* (knockout) embryos ((27) and figure S2A). We first analyzed if the commitment or delamination of LECs from the cardinal vein is affected in *Osr1^GCE/GCE^* embryos. Immunofluorescence of 100-µm thick sections from E11.5 *Osr1^+/+^* and *Osr1^GCE/GCE^*embryos showed no significant changes in LEC commitment or delamination into the surrounding mesenchyme together with normal blood vasculature structures (figure 2A and S2B). At E12.5, the first impairments in lymphatic structures were visualized by immunofluorescence in *Osr1^GCE/GCE^*embryos (figure 2B). Migration of LECs towards the dorsal region of the embryo appeared decreased and the PROX1+/VEGFR3+ vascular network in the primordial thoracic duct region showed reduced vascular branching (figure 2B). Whole-mount immunofluorescence of isolated back skin from E14.5 embryos showed markedly impaired migration of dermal LECs in the skin of E14.5 *Osr1^GCE/GCE^* embryos with significantly increased distances between growing fronts in both cervical and lumbar regions compared to *Osr1^+/+^* controls (figure 2C-E and S2C, D). In addition, lymphatics vessels showed a decrease in the numbers of branching points (figure 2C, F and G) and increased caliber (figure 2F, H) in E14.5 *Osr1^GCE/GCE^* skin. In line with the defects found in LEC migration, LECs in the migrating front of E14.5 *Osr1^GCE/GCE^* dermis presented a reduced number of filopodia per cell (figure S2E) suggesting an impaired ECM-LEC interaction. Of note, in the skin of E14.5 *Osr1^GCE/GCE^* embryos, blood vasculature showed no significant impairment in vessel thickness or branching point numbers (figure S2F).

**Figure 2.**
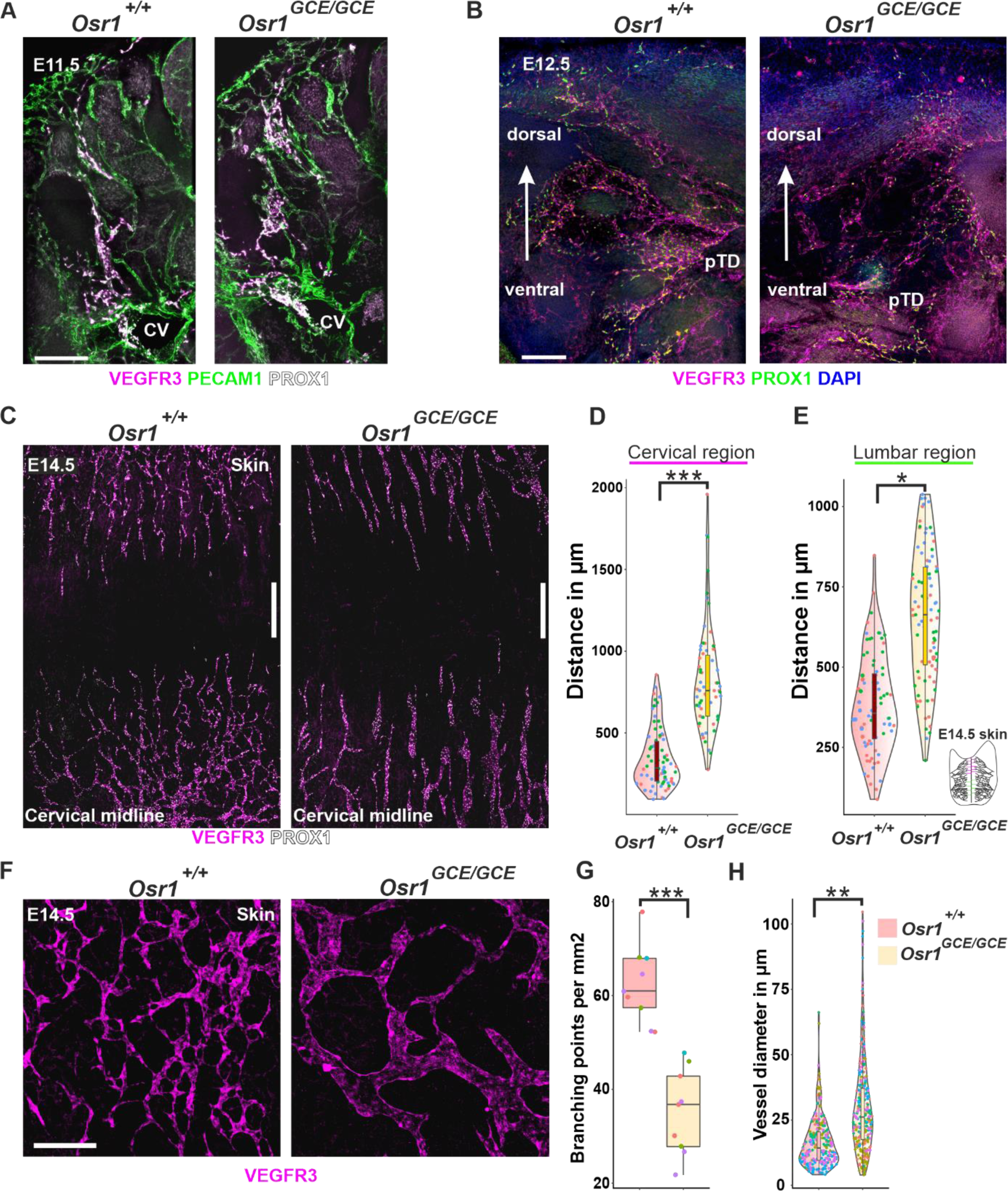
Lack of Osr1 in mesenchymal cells leads to lymphatic vasculature defects. **(A)** Maximal intensity projection images of Osr1+/+ and Osr1GCE/GCE E11.5 100 µm cross-sections after immunofluorescence. Immunolabeling for PROX1 and VEGFR3, or PECAM1, shows normal commitment and delamination of LECs and normal blood endothelial structures in the dorsal part of the cardinal vein. **(B)** At 12.5, maximal intensity projection images of 100 µm cross-sections reveal first impairments in lymphatic structures labeled for PROX1 and VEGFR3. **(C)** Whole-mount immunofluorescence of E14.5 Osr1+/+ and Osr1GCE/GCE skin samples for VEGFR3 and PROX1. **(D, E)** Quantification of the distance in µm between the tips of the migrating lymphatic vasculature front and the center of the avascular line in the skin in cervical and lumbar regions. n=3. **(F)** Representative micrographs of E14.5 Osr1+/+ and Osr1GCE/GCE skin labeled for VEGFR3. **(G)** Quantification of lymph vessel branching points per mm2 and **(H)** vessel diameter in E14.5 Osr1+/+and Osr1GCE/GCE skin labeled for VEGFR3. In **G**, n=4 and in **H** n=5. Measurements obtained from the same embryo are represented as dots with the same color. Representative immunofluorescence images have been captured from at least 3 different embryos. Scale bar represents in **A**, 200 µm, **B**, 100 µm, **C**, 500 µm and **F**, 200 µm. P values were obtained from student t tests. * represents p< 0.05, ** p< 0.01, *** p< 0.001. In **A**, CV represents cardinal vein. In **B**, pTD represents primordial thoracic duct.

Overall, this shows that the transcription factor *Osr1* expressed in mesenchymal cells is crucial for the formation of lymphatic vasculature, the loss of *Osr1* leads to severe impairments in LEC migration and lymphatic vasculature structures.

### Transcriptome analysis reveals impaired ECM interaction of LECs in E13.5 *Osr1-KO* embryos

At E13.5, *Osr1* is transcribed mainly in mesenchymal cell populations and is not detected in LECs (19, 28). To assess LEC defects in E13.5 *Osr1-*deficient embryos, we first isolated E13.5 LECs (PDPN+ PECAM1+) cells via fluorescence activated cell sorting (FACS) (figure 3A). Enrichment of LECs was confirmed via real-time (RT) qPCR analysis of key LEC markers *Flt4, Prox1* and *Ccl21* compared to FACS-isolated E13.5 blood endothelial cells (BECs) and mesenchymal *Osr1^GCE/+^* cells (figure 3B). Of note, relative percentages of BECs and LECs were not impaired in E13.5 *Osr1^GCE/GCE^*embryos (figure S3A) suggesting there was no general defect in LEC pool expansion and maintenance. Next, we performed transcriptome analysis of E13.5 LECs from *Osr1^controls^* and *Osr1^GCE/GCE^* embryos. This analysis revealed 1386 differentially expressed genes (DEG) with 877 down-, and 509 upregulated genes (figure 3C). We further characterized DEGs by performing gene ontology (GO) analysis. GO analysis for biological processes on all DEGs revealed an enrichment of terms associated with ECM organization and cell migration (figure 3D and S3B). ECM-associated terms were also identified by GO analysis using the Jensen compartment database (figure 3E), and the term ECM-receptor interaction was enriched after GO analysis using the Kyoto Encyclopedia of Genes and Genomes (KEGG) pathway database (figure 3F and S3C). Further GO analysis for biological processes by separating down and upregulated genes showed an enrichment of ECM terms in genes downregulated in LECs of *Osr1^GCE/GCE^*embryos (figure 3G). In line, GO analysis against the Jensen Compartment database by separating down and upregulated genes revealed genes associated with ECM-related terms as top ranked within downregulated genes (figure S3D). Within genes upregulated in LECs of E13.5 *Osr1^GCE/GCE^* embryos, GO analysis for biological processes revealed terms associated with vascular development ranking top (figure 3G). In line with this, selected genes known to have a positive effect in lymphatic vasculature formation (2, 8, 32–34) were found to be upregulated in LECs of E13.5 *Osr1^GCE/GCE^* embryos (figure 3H).

**Figure 3.**
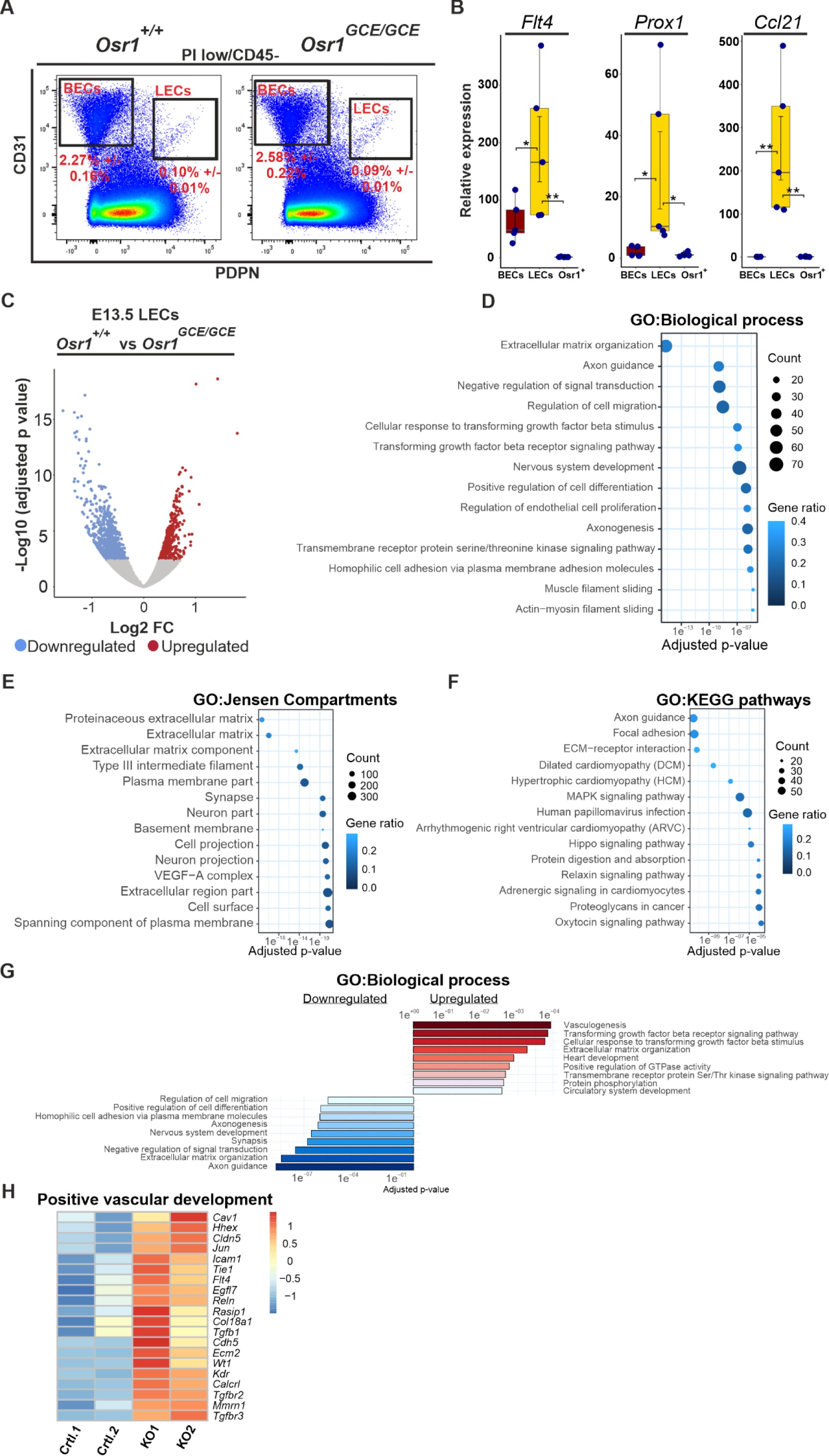
Transcriptional adaptations of LECs to an environment lacking Osr1. **(A)** FACs sorting strategy for isolating E13.5 LECs (CD45-PDPN+ CD31+) and BECs (CD45-PDPN-CD31+) from Osr1_controls_ (Osr1_+/+_ and Osr1_GCE/+_) and Osr1_GCE/GCE_ embryos. Percentage of cells +/-standard deviation of the mean is shown. n=9. **(B)** Boxplots from RT-qPCR analysis showing relative expression of Flt4, Prox1 and Ccl21 in E13.5 FACS isolated BECs, LECs and Osr1_GCE/+_cells. Relative expression was normalized to Osr1_GCE/+_ cells. n=5. **(C)** Volcano plot showing transcriptome analysis of E13.5 FACS sorted LECs from Osr1_controls_ and Osr1_GCE/GCE_ embryos showing upregulated (red) and downregulated (blue) genes identified by an absolute log2 fold change above 0.3 and an adjusted p-value below 0.05. **(D-F)** Dot plot depiction of GO analysis for biological processes, Jensen Compartments and KEGG pathways using all deregulated genes; top 14 terms ranked by their adjusted p-value are shown. Count represents number of genes in the term and Gene ratio the percentage of significant genes over the total genes in a given term. **(G)** Bar plot representation of GO analysis for biological processes performed in up (red) or downregulated genes (blue). Terms were ranked by their adjusted p-value. **(H)** Heatmap depiction of TPM values for selected genes positively involved in lymphatic vessel formation. Raw scaled normalization is represented at the right. In **B**, p values were obtained from one-way ANOVA with Dunnett’s multiple comparisons. Error bar represents s.e.m. and * p< 0.05 and ** p< 0.01.

In summary, lack of *Osr1* in mesenchymal cells leads to a series of transcriptional adaptations in embryonic LECs. Mesenchymal cell impairment leads to decreased expression of ECM and ECM-receptor interaction genes and downregulation of genes involved in cell migration in LECs of *Osr1^GCE/GCE^* embryos. Finally, E13.5 *Osr1^GCE/GCE^* LECs activate genes positively involved in lymphatic vessel formation suggesting a compensatory mechanism.

### Transcriptome analysis of mesenchymal Osr1+ cell - LEC interactions via the ECM

RNA-seq analysis revealed that LECs in *Osr1^GCE/GCE^* embryos have a deregulated transcriptional signature of ECM organization and ECM-receptor interaction with many genes related to these terms being downregulated (figure 3 and S3). To address this from the perspective of Osr1+ mesenchymal cells, we used our previously published (28) RNA-seq dataset of Osr1+ cells from E13.5 *Osr1^GCE/+^* vs. *Osr1^GCE/GCE^* embryos. To specifically select for dermal fibroblast-expressed genes, we intersected the 511 E13.5 *Osr1^GCE/GCE^* DEGs with the 976 genes found to be highly abundant in the dermal cluster of E13.5 *Osr1* cells identified by scRNA-seq analysis (19), given that this cluster was enriched for key embryonic skin dermal fibroblast population markers (35, 36) (figure 4A). Amongst the genes deregulated in E13.5 *Osr1^GCE/GCE^*cells, we identified 50 deregulated “dermal” genes (figure 4B). We subjected this set of genes to GO analysis and found the terms small leucine-rich proteoglycan (SLRP) molecules, NCAM1 interactions, and TGF-beta regulation of extracellular matrix as significant terms downregulated in this intersection (figure 4C). GO analysis for molecular functions of only upregulated genes in this intersection highlighted metalloendopeptidase and metallopeptidase activity as the only significantly enriched terms after the analysis (figure 4D). In line with the ECM defects observed in embryos lacking *Osr1* in other tissues (28, 29), this analysis suggested that in dermal E13.5 Osr1-deficient cells, ECM production and organization may be also impaired.

**Figure 4.**
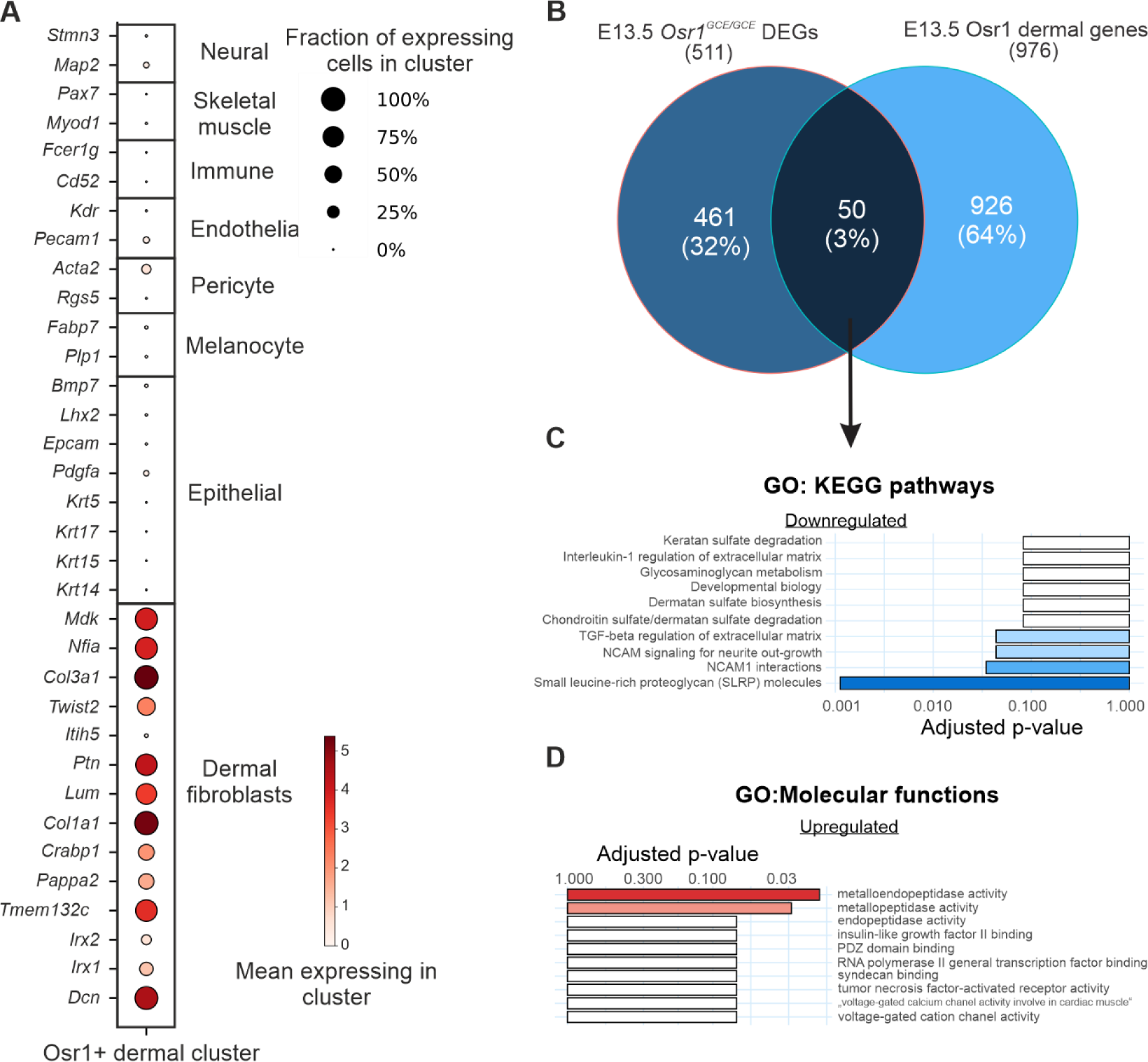
LECs-Osr1 interactions in the dermis. **(A)** Expression of key genes for the main cell populations of embryonic skin tissues (37) in the E13.5 Osr1+ dermal cell cluster identified by single cell RNA-Seq (19). **(B)** Venn diagram intersecting deregulated genes obtained after RNA-seq analysis of E13.5 Osr1_GCE/+_ vs Osr1_GCE/GCE_ cells and genes characterizing the E13.5 Osr1 dermal cluster identified by scRNA-seq analysis. **(C, D)** Bar plot representation of GO analysis using 50 genes deregulated in Osr1_GCE/GCE_ cells and enriched in E13.5 dermal cluster for KEGG pathways in downregulated genes (blue) and for molecular functions in upregulated genes (red). Terms were ranked by their adjusted p-value.

### Impaired ECM scaffold in the dermis of *Osr1* deficient embryos

Transcriptome analyses of E13.5 Osr1+ cells and LECs suggested defective ECM organization in the developing lymph vasculature of *Osr1^GCE/GCE^*embryos. We therefore characterized the ECM in direct contact with the dermal vasculature. Dermal fibroblasts are the primary source of ECM scaffold components such as *Col12a1*, *Col1a1*, *Fn1*, *Col3a1*, or SLRPs as *Dcn* or *Lum*; of note, *Osr1* expression is enriched in dermal fibroblasts (37) (figure S4A). Whole-mount immunofluorescence of E14.5 skin using antibodies against the ECM proteins COL12A1, COL6 and TNC showed an impaired ECM scaffold in direct contact with lymphatic vasculature showing frayed fiber organization and increased TNC expression in E14.5 *Osr1^GCE/GCE^* embryos (figure 5A-C). Of note, ECM-organization defects were observed in ventral regions as well as in the region of the dorsal migrating front of LECs (figure S4B). Despite normal formation of dermal blood vessels observed in E14.5 *Osr1^GCE/GCE^* embryos (figure S2F), closer appreciation of the basal lamina in dermal blood capillaries showed a reduced accumulation of COL1 protein together with an impaired COL1 deposition (figure 5D and S4C).

**Figure 5.**
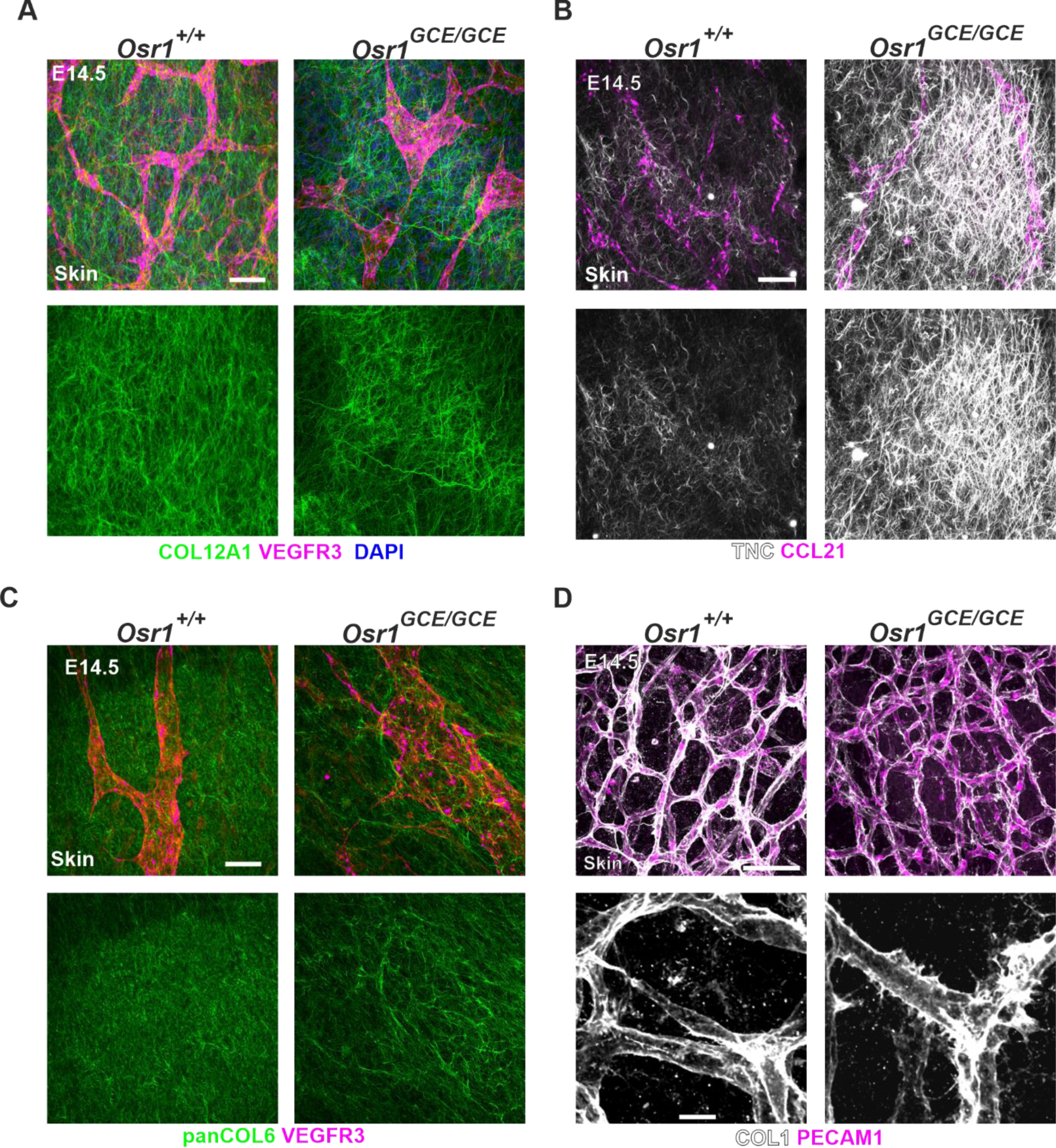
Impaired dermal ECM in E14.5 Osr1_GCE/GCE_ embryos. **(A-C)** Representative micrographs of E14.5 skin whole-mount immunofluorescence showing ECM impairments in Osr1_GCE/GCE_ embryos. ECM was labeled for COL12A1, TNC and COLVI. Lymphatic vasculature is labeled for VEGFR3 and CCL21. **(D)** Representative micrographs of E14.5 skin whole-mount immunofluorescence showing defects in the basal lamina of blood vessels. Endothelial cells are labeled for PECAM1 and basal lamina for COL1. Representative immunofluorescence images have been captured from at least 3 different embryos. Scale bar represents in **A-D**, 50 µm and in **D below**, 20 µm.

### Mesenchymal Osr1+ cells are a source of *Vegfc* and control LEC proliferation via a proactive-ECM

Activation of the VEGFC/VEGFR3 signaling pathway is essential for lymphatic vessel formation driving LEC delamination from the cardinal vein and LEC proliferation (10, 38–41). Available sc-RNAseq data for the stages E9.5-E13.5 (42) showed that mesenchymal cells and endothelial cells were the major source of *Vegfc* expression (figure S5A). We confirmed that E13.5 Osr1+ cells showed high expression of *Vegfc* together with endothelial cells by comparing *Vegfc* transcriptional expression in Osr1+ cells, BECs and LECs separated by FACS (figure 6A). At E11.5, *Vegfc* expression in BECs was higher compared to mesenchymal Osr1+ cells, however at E13.5 Osr1+ cells were the major cell type expressing *Vegfc* (figure S5B). Transcriptome analysis of E13.5 *Osr1^GCE/+^*and E13.5 *Osr1^GCE/GCE^* mesenchymal cells revealed a decrease in *Vegfc* transcripts in cells lacking *Osr1* (28). We confirmed *Vegfc* transcript downregulation in E13.5 *Osr1^GCE/GCE^* mesenchymal cells, whereas at E11.5, *Osr1^GCE/GCE^* mesenchymal cells produced similar *Vegfc* transcripts compared to controls (figure 6B). Contradictory to the notion that reduced *Vegfc* in mesenchymal cells may be responsible for the lymphatic vessel defects observed in *Osr1* deficient embryos, we observed an increase in *Flt4* expression in LECs of E13.5 *Osr1^GCE/GCE^*embryos (figure 3F). Besides, decreased *Vegfc* expression in mesenchymal cells did not implicate a reduction at the transcriptional level of genes involved in the VEGFR3-signaling axis in E13.5 LECs from *Osr1^GCE/GCE^* embryos. Indeed, *Flt4, Hhex* and other downstream targets of the signaling pathway such as *Maf*, or *Egr1* (15, 32, 43, 44) were upregulated in LECs from E13.5 *Osr1^GCE/GCE^* embryos (figure 6C). Since BECs are also a source of *Vegfc*, we analyzed *Vegfc* expression in E13.5 *Osr1^GCE/GCE^* embryos via RT-qPCR analysis of FACS isolated BECs and did not observe a change in *Vegfc* expression (figure S5C). Of note, available transcriptome analysis of E14.5 skin populations confirms fibroblast and melanocytes as the main cell types expressing *Vegfc* (45) (figure S5D).

**Figure 6.**
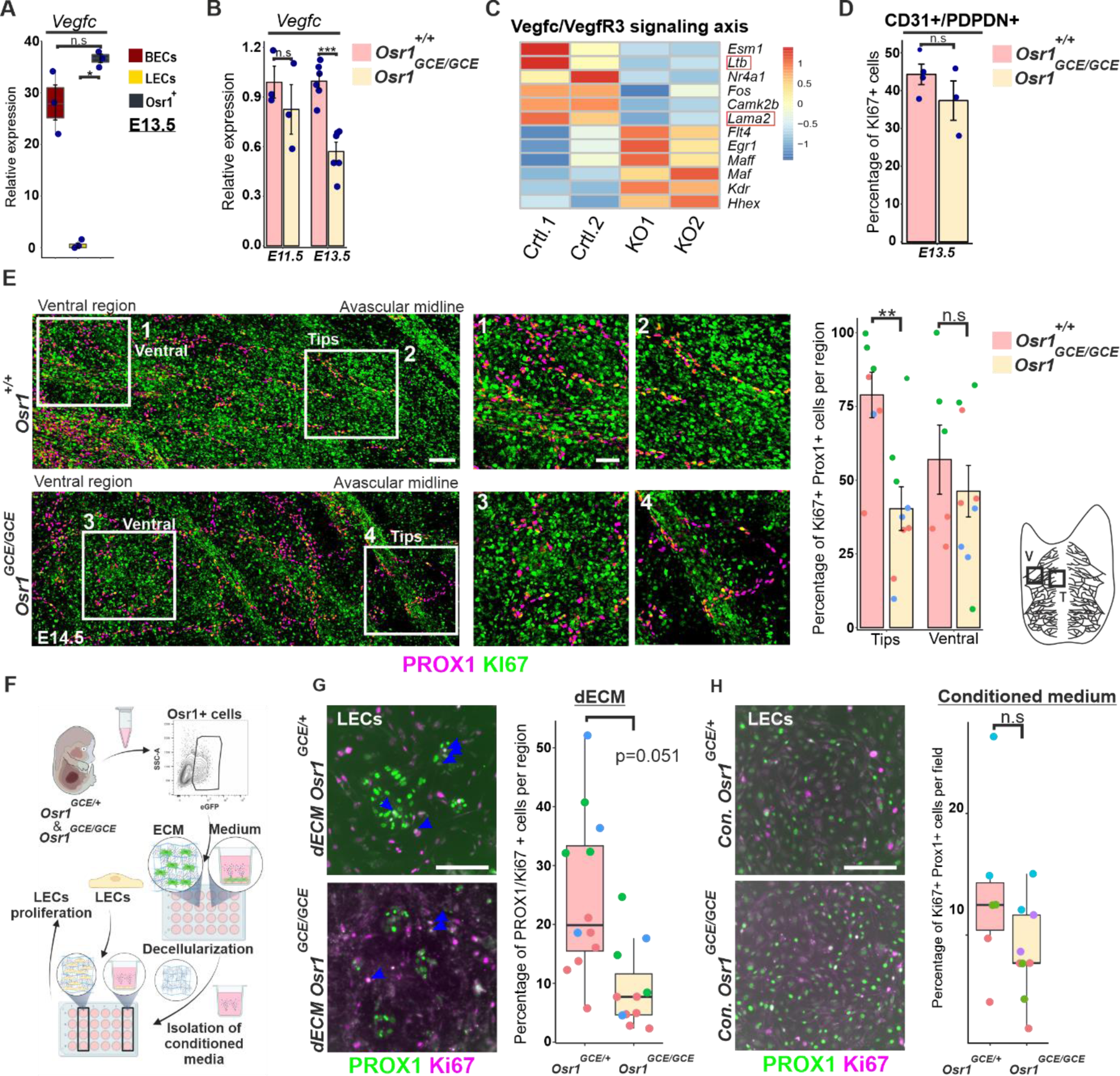
Mesenchymal Osr1+ cells promote LEC proliferation at the migrating front via the ECM. **(A)** Relative expression of Vegfc in E13.5 Osr1_GCE/+_ cells, BECs and LECs isolated by FACS. n=3. **(B)** RTqPCR analysis showing Vegfc relative expression in E11.5 and E13.5 FACS isolated Osr1_GCE/+_and Osr1_GCE/GCE_ cells. In E11.5, n=3 and in E13.5 n=6. **(C)** Heatmap depiction of TPM values for selected genes involved in the VEGFC/VEGFR3 signaling pathway. Raw normalized scale is represented at the right. Genes surrounded by a red square are not in agreement with upregulation of VEGFC/VEGFR3 signaling pathway described by others. **(D)** Quantification of Ki67+/PROX1+ LECs from E13.5 Osr1_GCE/+_and Osr1_GCE/GCE_ embryos isolated by FACS that shows no differences in proliferation. n=3. **(E)** Representative micrographs of E14.5 skin whole-mount immunofluorescence showing Ki67+ proliferative LECs labeled by PROX1. Quantification of PROX1+/Ki67+ cells in ventral (V) and migration front (T) regions is shown at the right together with a schematic representation of skin lymphatic vessels and regions measured. n=4**. (F)** Schematic representation of experimental workflow for decellularized (d)ECM and conditioned media production from E13.5 FACS isolated Osr1_GCE/+_and Osr1_GCE/GCE_ cells. Subsequent pdLECs culture was performed on dECM or using conditioned media. **(G)** Immunofluorescence of pdLECs cultured on dECM for 48 hours. LECs are labeled for PROX1 and Ki67 for proliferation. Quantification of proliferative LECs is shown at the right. n=2-3. **(H)** Immunofluorescence of pdLECs cultured for 24 hours in conditioned media coming from E13.5 FACS isolated Osr1_GCE/+_and Osr1_GCE/GCE_ cells. LECs are labeled for PROX1 and Ki67 for proliferation. Quantification of proliferative LECs is shown at the right. n=3-4. Measurements obtained from the same embryo are represented as dots with the same color. Representative immunofluorescence images have been captured from at least 3 different embryos. Scale bar represents in **E**, 100 µm and 50 µm (1–4), in **G, H**, 200 µm. In **B**, p values were obtained from one-way ANOVA with Dunnett’s multiple comparisons. In **A, D, E, G** and **H**, p values were obtained from student t tests. Error bar represents s.e.m. and * represents p< 0.05, ** p< 0.01, *** p< 0.001 and n.s not significant.

E13.5 LECs isolated by FACS from E13.5 whole *Osr1^+/+^* and *Osr1^GCE/GCE^* embryos and cultured *in vitro* did not show defects in cell proliferation (figure 6D). Interestingly, we detected using whole-mount immunofluorescence a heterogeneous proliferation of LECs depending on their distribution in the skin. LECs in the migrating front show higher proliferation quantified by PROX1/Ki67 co-staining, whereas in the ventral side, LECs of the established lymphatic vasculature were less proliferative (figure 6E). Quantification of E14.5 LECs proliferation showed a reduced proliferation in E14.5 *Osr1^GCE/GCE^* embryos specifically in the migrating front, while LECs in the ventral lymphatic vasculature (figure 6E) showed similar proliferation in *Osr1*-deficient and control embryos.

In order to untangle the mechanism used by Osr1+ mesenchymal cells to control LEC proliferation, we aimed to separate the effects coming from signaling molecules secreted by Osr1+ cells and the effects of the ECM scaffold produced by Osr1+ cells. For this propose, we isolated E13.5 *Osr1^GCE/+^* and *Osr1^GCE/GCE^* cells via FACS and let them produce either an ECM scaffold or a conditioned medium (figure 6F). Next, we isolated dermal LECs from the tail (tdLECs) of adult mice (figure S5E, F) and quantified the effects of E13.5 *Osr1^GCE/+^* and *Osr1^GCE/GCE^*decellularized ECM (dECM) or Osr1+ cell-conditioned media on tdLECs proliferation (figure 6F). LECs cultured for 48h on E13.5 *Osr1^GCE/GCE^* dECM showed reduced proliferation quantified by PROX1+/Ki67+ co-staining as compared to dECM produced by E13.5 *Osr1^GCE/+^* cells (figure 6G). Conversely, conditioned medium produced by E13.5 *Osr1^GCE/GCE^* cells did not significantly reduce LECs proliferation as compared to E13.5 *Osr1^GCE/+^*conditioned medium (figure 6H).

In conclusion, while mesenchymal Osr1+ cells are an important source of *Vegfc* in the mouse, decreased *Vegfc* in *Osr1^GCE/GCE^*cells did not lead to decreased VEGFR3 downstream target expression in LECs. Instead, a defective ECM secreted by *Osr1^GCE/GCE^* cells affects LEC proliferation *in vitro* in line with decreased LEC proliferation in E14.5 *Osr1^GCE/GCE^* embryos at the migrating front of the growing lymph vasculature.

### Mesenchymal Osr1+ cells provide beneficial guidance for LEC migration

In line with a reduced LEC migration observed in the skin of E14.5 *Osr1^GCE/GCE^* embryos, cell migration is one of the most enriched terms in the GO analysis for biological processes in E13.5 LEC RNA-seq data, where most deregulated genes in that term appeared downregulated (figure 7A). To clarify if *Osr1* controls the production of signaling molecules expressed by mesenchymal cells acting on LECs and controlling their migration, we assessed LEC migration *in vitro* by performing scratch assays in cultures of tdLECs isolated from the dermis of adult tail tissues supplemented with conditioned media produced by E13.5 *Osr1^GCE/+^* and *Osr1^GCE/GCE^* cells. TdLECs cultured for 24 hours in conditioned medium from E13.5 *Osr1^GCE/GCE^*cells migrated slower into the acellular space compared to LECs cultured in conditioned medium from E13.5 *Osr1^GCE/+^* control cells (figure 7B). To evaluate putative interactions between E13.5 Osr1+ cells and LECs, we analyzed ligand-receptor interactions using all deregulated genes found in E13.5 *Osr1^GCE/GCE^*cells (511 genes) and in E13.5 LECs (1386 genes) that matched with the interacting pairs defined in (46) (figure 7C). Within the interactions of deregulated ligands in Osr1+ cells and deregulated LEC receptors, we found that the Osr1-LEC ligand-receptor pairs TNC-EGFR/ITGA9/ITGAV represented interactions of genes found upregulated in both cell types in line with increased TNC protein abundance in *Osr1^GCE/GCE^*skin. By contrast, the ligand-receptor pairs, COL3A1-DDR1*/*2 and CXCL12-CXCR4, depicted highly expressed ligands by Osr1+ cells (*Col3a1* and *Cxcl12*) found downregulated in E13.5 *Osr1^GCE/GCE^* cells and their interaction partners (*Ddr1*/*2* and *Cxcr4*) downregulated in LECs. The chemokine *cxcl12* controls LEC migration in zebra fish development and in newborn mice via its receptors *cxcr4* (16, 18). Therefore, we asked whether *Cxcl12* might have a similar function in embryonic dermal lymphatic vasculature formation in mouse embryos. First, we assessed *Cxcl12* expression in E13.5 Osr1+ cells, BECs and LECs and observed that mesenchymal Osr1+ cells appear as the main source of *Cxcl12* expression (figure 7D). Next, we used the *Cxcr4^KO/KO^* line (47) and performed whole-mount immunofluorescence for lymphatic endothelial cells markers. At E14.5 LECs showed impaired migration to the dorsal midline in E14.5 *Cxcr4^KO/KO^*embryos with a concomitant increase in lymphatic vessel caliber and reduced arborization (figure 7E and S7A), similar to E14.5 *Osr1^GCE/GCE^* embryos. In line with the function assigned to *cxcl12* in zebra fish (16), LECs in *Cxcr4^KO/KO^* embryos did not show a reduced proliferation at the tip of the migrating front (figure 7F). This suggest that in the mouse CXCL12/CXCR4 signaling is required for LEC dorsal migration downstream of mesenchymal *Osr1*.

**Figure 7.**
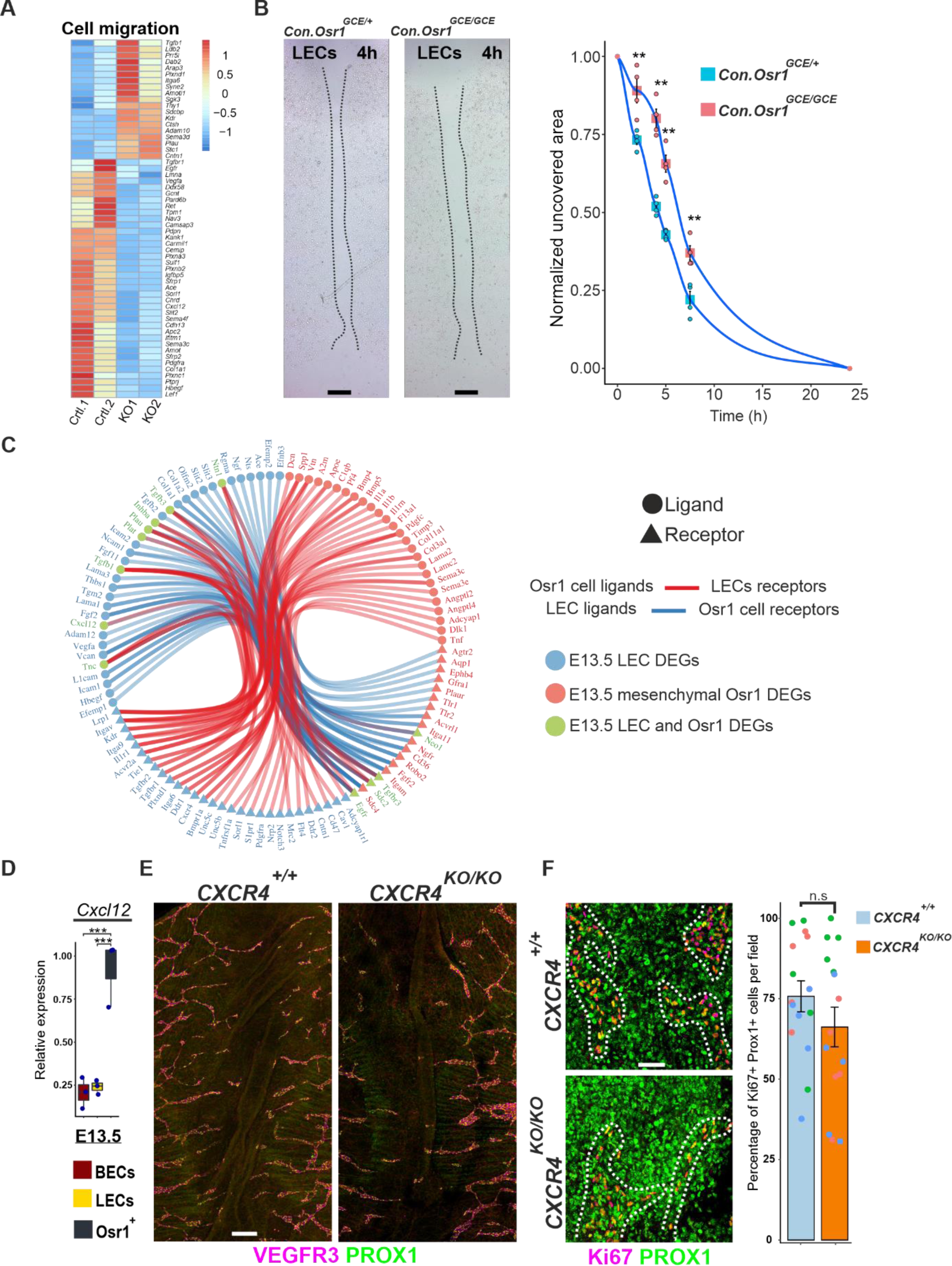
Mesenchymal Osr1+ cells promote LEC migration. **(A)** Heatmap depiction of TPM values of genes belonging to the GO term cell migration. Raw scaled normalization is represented at the right. **(B)** Migration assay using tdLECs under conditioned media produced by E13.5 Osr1_GCE/+_ and Osr1_GCE/GCE_ cells. Representative images after 4h of culture shown left, quantification of wound closure at 0, 2, 4, 5, 7.5 and 24h after induction with conditioned media shown right. N=4. **(C)** Ligand-receptor pair analysis using DEGs of E13.5 Osr1_GCE/GCE_ cells and LECs from E13.5 Osr1_GCE/GCE_ embryos. **(D)** RT-qPCR analysis showing Cxcl12 relative expression in E13.5 FACS isolated Osr1_GCE/+_ BECs and LECs isolated by FACS. N=3. **(E)** E14.5 skin whole-mount immunofluorescence of Cxcr4_+/+_ and Cxcr4_KO/KO_ embryos showing lymphatic vessel impairments. Lymphatic vasculature is labeled for VEGFR3. **(F)** Representative micrographs of E14.5 skin whole-mount immunofluorescence showing normal proliferation of LECs in the migrating front from Cxcr4_KO/KO_ embryos. LECs are labeled for PROX1 and proliferation measured by Ki67. Quantification of Ki67+/PROX1+ cells per region is shown at the right. N=4. Representative immunofluorescence images have been captured from at least 3 different embryos. Scale bar represents in **B**, 500 µm, in **E**, 200 µm and in **F**, 50 µm. In **D**, p values were obtained from one-way ANOVA with Dunnett’s multiple comparisons. In **B, F**, p values were obtained from student t tests. Error bar represents s.e.m. and * represents p< 0.05, ** p< 0.01, *** p< 0.001 and n.s not significant.

## Discussion

The formation of lymphatic vasculature during development is crucial for body fluid homeostasis and therefore for animal survival. Although several steps in lymphatic vessel development have been well characterized, the role of mesenchymal cells in lymphatic vessel development remained understudied. We have shown that mesenchymal Osr1+ cells accompany the early migration path of LECs from the cardinal vein to peripheral tissues including the mesenchyme that surround the primordial thoracic duct, and Osr1+ cells remain in close association with lymphatic vasculature in several vascular beds. Lineage tracing experiments suggest that mesenchymal Osr1+ cells appear in the mesenchyme preceding the LEC migration front in the dermis of the embryo. Osr1+ descendants also contribute to mural cells of arteries and veins; it remains to be elucidated if Osr1+ cells constitute a population of mural cells for mature lymphatic vessels.

Interestingly, in the medulla of lymph nodes, *Osr1* expression remains active in close association to blood and lymphatic vessels, and *Osr1* and *Vegfc/Vegfa* expression are both found in lymph node mesenchymal stromal subpopulations (data not shown, Cyster Lab Shyni Server).

Reminiscent of the activation of *Osr1* expression in skeletal muscle mesenchymal cells triggered by acute muscle injury (29), we observed transcriptional reactivation of *Osr1* in the dermis of the ear after trauma. This indicates that *Osr1* reactivation may represent a general mechanism employ by mesenchymal cells as a trauma response, possibly to induce a pro-remodeling phenotype.

Lack of *Osr1* in mesenchymal cells in *Osr1^GCE/GCE^*embryos led to impairment of lymphatic vasculature formation indicating that Osr1+ mesenchymal cells are a new important player controlling LEC behavior in a non-cell autonomous fashion. In line with the function assigned to mural cells in zebra fish controlling LEC migration and survival (16), we observed LEC migration defects in *Osr1* deficient embryos and a reduced dermal LEC proliferation specifically at the tips of the migrating front. Conversely, blood vessel morphology and pattern was not significantly impaired in *Osr1*-deficient embryos. Of note, the collagen-rich basal lamina of capillaries in the skin of E14.5 *Osr1^GCE/GCE^* embryos shows defects in ECM deposition, hinting at a possible function of Osr1+ cells in blood vessel stability, which remains to be explored. The differential impact of *Osr1* deficiency on blood and lymphatic vasculature formation is in agreement with the earlier formation of the blood vascular plexus in the embryo (48) and a later appearance of Osr1+ mesenchymal cells in close proximity to blood vasculature (26, 49).

We found that Osr1+ cells are a source of *Vegfc* in embryonic tissues. Interestingly, *Vegfc* expression at early stages of development was higher in endothelial cells (42) while at E13.5 Osr1+ mesenchymal cells and BECs expressed similar amounts of *Vegfc*. Despite *Vegfc* downregulation observed in E13.5 *Osr1^GCE/GCE^* mesenchymal cells, the LEC transcriptome data suggested that embryonic E13.5 LECs show even increased activation of the VEGFC/VEGFR3 signaling pathway, at least when assessed at the transcriptional level. This might represent a compensatory mechanism in line with the transcriptional upregulation of genes positively involved in vascular development observed in GO analysis of E13.5 LECs from *Osr1^GCE/GCE^* embryos. An alternative hypothesis explaining the upregulation of genes involved in the VEGFC/VEGFR3 axis in LECs includes an increase in VGEFC final active form resulted from an increase in protease activity in the dermis of *Osr1* deficient embryos. Collectively, our results suggest that the dysfunctional LEC behavior observed in *Osr1^GCE/GCE^*embryos is not caused by a perturbed VEGFC/VEGFR3 signaling axis.

In the embryonic dermis, fibroblasts are the main producers of ECM components and therefore key players in the formation of the ECM scaffold embedding dermal blood and lymphatic vasculature (45). Lack of *Osr1* leads to a severely disorganized ECM scaffold in the dermis of E14.5 embryos paralleling our previous observations in skeletal muscle (28). In addition, the most prominent terms after GO analysis of deregulated genes in LECs of E13.5 *Osr1^GCE/GCE^* embryos are related to ECM and ECM-interaction genes, suggesting that LECs themselves react to the altered ECM produced by Osr1-deficient mesenchymal cells. In line, decellularized ECM, but not conditioned medium, produced by E13.5 *Osr1^GCE/GCE^* cells *in vitro* failed to properly sustain LEC proliferation. This suggests that the reduced proliferation of LECs at the tips of the migrating zone in the dorsal skin of *Osr1^GCE/GCE^* embryos was caused by aberrant ECM deposition from *Osr1*-expressing mesenchymal cells. This agrees with previous reports showing that ECM composition and stiffness can modulate BEC (50–52) and LEC (24) behavior.

Moreover, ECM and genes involved in endothelial ECM interactions play a role in controlling endothelial migration and stability (53–55). Interestingly, *Amot* and genes defined to interact directly with AMOT proteins such as *Kank1*, *Kank3* and *Flnc* (54) are downregulated in LECs of E13.5 *Osr1^GCE/GCE^* embryos (figure S7B). We also observed a reduced formation of filopodia at the tips of migrating LECs in E14.5 *Osr1^GCE/GCE^* embryos suggesting altogether a defective ECM-LEC interaction that may affect also LEC migration.

In addition to producing the bulk of ECM, mesenchymal cells express signaling molecules that control LEC migration (16). Conditioned media experiments suggested that Osr1+ mesenchymal cells produce signaling molecules necessary for LEC migration, and transcriptome-based Osr1 cell – LEC interaction analysis highlighted the CXCL12/CXCR4 axis as possible mechanism. Of note, *Cxcl12* expression is directly regulated by *Osr1* and Cxcl12 is highly expressed by Osr1+ cells (28). In support of this idea, *CXCR4^KO/KO^* embryos displayed similar defects in lymphatic vessel formation as *Osr1^GCE/GCE^* embryos. However, in the dermis of E14.5 *CXCR4^KO/KO^* embryos, LEC proliferation at the tip of the migrating front was not affected. In agreement, cxcr4 inhibition in zebra fish affected mainly LEC migration and not their proliferation (16). This suggests that the CXCL12/CXCR4 signaling pathway may be responsible for LEC migration during lymphatics development in the mouse as in zebra fish, while LEC proliferation, specifically at the migration front, is controlled by the ECM.

In summary our data show that mesenchymal Osr1+ cells play a fundamental role in LEC migration, proliferation and lymphatic vasculature assembly. Hereby *Osr1* is a key player controlling in a bimodal manner the production of critical ECM scaffold components and signaling ligands to provide a microenvironment necessary for lymphatic vessel formation. This parallels the mode of action applied by Osr1+ cells in the developmental formation and adult regeneration of skeletal muscle (28, 30) and the developmental formation of lymph nodes (19), and thus suggests an overarching mechanism by which mesenchymal cells control organ formation with the transcription factor *Osr1* at a key nexus.

## Acknowledgements

This work was funded by the Deutsche Forschungsgemeinschaft (DFG; grant VA 1272/1-1) and the Freie Universität Berlin. René Hägerling was supported in part by the Berlin Institute of Health (BIH) and by grants from the Lymphatic Malformation Institute and European Union (ERC, PREVENT, 101078827). We gratefully acknowledge Andrew P. McMahon (Keck School of Medicine of USC, USA) and Andreas Kispert (Hannover Medical School, Germany) for providing *Osr1^GCE^* and *R26^mTmG^*mouse lines. We thank Carmen Birchmeier and Ines Lahmann (Max Delbrück Center for Molecular Medicine, Berlin, Germany) for providing *Cxcr4*^+/-^ mice. We thank Uta Marchfelder and Erwin Weiß (Max Planck Institute for Molecular Genetics, Germany) for flow cytometry support. We thank Stefan Mundlos, Nobert Brieske and Thorsten Mielke for their support (Max Planck Institute for Molecular Genetics, Berlin, Germany). We are grateful to Olaf Penack (Charité, Berlin, Germany) for his support.

## Author contributions

Conceptualization P.V-G, M.O, R.H. and S.S. P.V-G, M.K., N.H and G.K. performed data collection. P.V-G and M.O. performed transcriptome analysis. C.G-T supervised FACS experiments. B.T supervised sequencing experiments. Formal analysis and interpretation were performed by P.V-G, M.O., R.H. and S.S. P.V.G and S.S wrote the manuscript.

All authors critically reviewed and approved the final version of the manuscript.

